# Weak predation strength promotes stable coexistence of predators and prey in the same chain and across chains

**DOI:** 10.1101/2020.02.09.940874

**Authors:** Lin Wang, Yan-Ping Liu, Rui-Wu Wang

## Abstract

The mechanisms of species coexistence make ecologists fascinated, although theoretical work show that omnivory can promote coexistence of species and food web stability, it is still a lack of the general mechanisms for species coexistence in the real food webs, and is unknown how omnivory affects the interactions between competitor and predator. In this work, we first establish an omnivorous food web model with a competitor based on two natural ecosystems (the plankton community and fig-fig wasp system). We analyze the changes of both food web structure and stability under the different resource levels and predation preference of the generalist/top predator. The results of model analyses show that weak predation strength can promote stable coexistence of predators and prey. Moreover, evolutionary trend of food web structure changes with the relative predation strength is more diversity than the relative competition strength, and an integration of both omnivory, increased competition, top-down control and bottom-up control can promote species diversity and food web stability. Our theoretical predictions are consistent with empirical data in the plankton community: the lower concentration of nutrient results in a more stable population dynamics. Our theoretical work could enrich the general omnivorous theory on species coexistence and system stability in the real food webs.

## 1. Introduction

Biodiversity in natural communities has both puzzled and fascinated ecologists [Wilson, 1992; Neutel et al., 2002]. Science Magazine celebrates the journal’s 125th anniversary with a look forward–at the most compelling puzzles and questions facing scientists, and one of the 25 most key questions is: what determines species diversity [Kennedy & Norman, 2005]? Species diversity in ecological communities has stimulated a century of study on the mechanisms that maintain coexistence of species [Lotka, 1925; Volterra, 1926; Gravel et al., 2011; Levine et al., 2017]. Understanding species coexistence is not only an intellectual challenge, but can also help restrain biological invasions [MacDougall et al., 2009] and protect rare species [DeCesare et al., 2010]. Until now, many mechanisms of species coexistence have been extensively discussed, such as natural enemies [Janzen, 1970; Connell, 1971], resource partitioning [Tilman, 1982] and responses to spatiotemporal variation [Chesson, 2000]. However, most of these work are based on a single theoretical perspective, there is still a lack of the general mechanisms of species coexistence in the real ecosystems.

An ecological community has usually been treated as a network of species connected by different kinds of interactions, such as predation, parasitism, mutualism and competition. Indeed, linking these interaction networks to community structure and stability may be a key to understanding the maintenance of ecological communities [Ushio et al., 2018]. As a general trophic structure of ecosystem, omnivorous link (connections across trophic levels) has attracted an enormous amount of work, such as theoretical models [McCann & Hastings, 1997; Fussmann & Heber, 2002; Tanabe & Namba, 2005; Sieber & Hilker, 2011] and empirical analyses [Snyder & Ives, 2003; Novak, 2013; Wilken et al., 2014]. Although early work show that omnivory can promote species coexistence and food web stability [McCann & Hastings, 1997; Fussmann & Heber, 2002], most of them are based on a tri-trophic interactions among prey, predator and superpredator: the superpredator also feeds on prey [McCann & Hastings, 1997; Tanabe & Namba, 2005; Sieber & Hilker, 2011]. The system is the simplest model with omnivory, which makes it manageable to do the theoretical analysis. However, it could be unsuitable to extend corresponding conclusions to the natural ecosystems.

In both terrestrial and aquatic ecosystems, for predicting species coexistence, the simplest structure of the omnivorous food webs will be affected by other modules, such as competitor [Compton & Robertson, 1988; Persson et al., 2001]. For example, in the fig-fig wasp system [Compton & Robertson, 1988], both ant, parasitoid and pollinator constitute the simplest omnivorous food web structure, while non-pollinator (seed predator) will compete with the pollinator for seeds. Moreover, the ant also feeds on the non-pollinator to protect the fig-fig wasp mutualism. The pollinator can be regarded as a mutualist due to it pollens the fig tree. In the plankton community [Persson et al., 2001; Benincà et al., 2008], both zooplankton (daphnia, small grazers), heterotrophic nanoflagellates and bacteria constitute the simplest omnivorous food web structure, while algae (edible algae, filamentous algae) will compete with bacteria for nutrients, and zooplankton also feeds on algae. The bacteria can be treated as a mutualist because of its decomposition can produce nutrients. Both of the two natural ecosystems consist of four modules: mutualist, competitor, specialist predator and generalist predator.

The interactions between predator and competitor were extensively discussed [Hatcher et al., 2006; Chesson & Kuang, 2008], both the theoretical models [Fussmann & Heber, 2002; Revilla & Weissing, 2008] and experimental work [Passarge et al., 2006; Foster & Bell, 2012] found that competition can promote species coexistence. However, it is still unknown how the omnivory affects the interactions between competitor and predator in the natural ecosystems.

In this work, we first establish an omnivorous food web model among mutualist, competitor, specialist predator and generalist predator based on two natural ecosystems. Then, we analyze how predation preferences of the generalist predator affect the interactions between competitor and specialist predator under the different resource levels. The results of model analyses show that weak predation strength can promote stable coexistence of predators and prey. Moreover, evolutionary trend of food web structure changes with the relative predation strength is more diversity than the relative competition strength, and an integration of both omnivory, increased competition, top-down control and bottom-up control can promote species diversity and food web stability. Finally, we have a contrastive analysis on the theoretical prediction and empirical data. Our theoretical work could enrich the general omnivorous theory on species coexistence and system stability in the real food webs.

## 2. Methods

### 2.1. Study system

In order to study the general mechanisms for species coexistence in the natural ecological systems, we first elicit the model framework based on both terrestrial and aquatic ecosystems (Fig. 1).

**Fig. 1.**
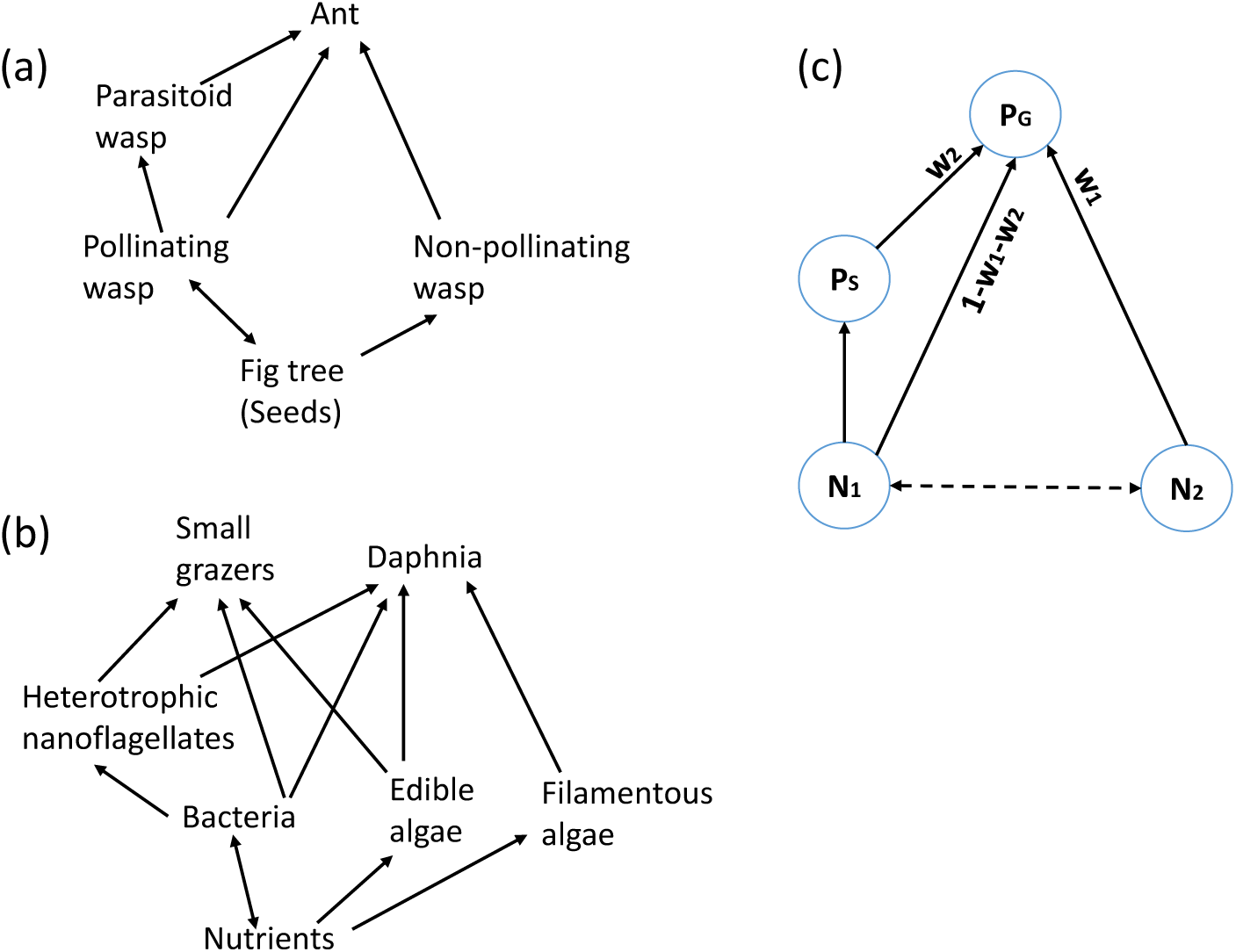
A graph of relationships among generalist predators (*P*_*G*_: Ant; Daphnia, Small grazers), specialist predators (*P*_*S*_ : Parasitoid wasp; Heterotrophic nanoflagellates), mutualist (*N*_1_: Pollinating wasp; Bacteria), competitors (*N*_2_: Non-pollinating wasp; Edible algae, Filamentous algae). (a) The fig-fig wasp mutualistic system (Compton & Robertson, 1988); (b) Plankton community (Persson et al., 2001; Benincà et al., 2008); (c) A model framework of the omnivorous food web model among mutualist, competitor, specialist predator and generalist predator. Both *w*_1_ and *w*_2_ are the predation preference coefficients. The solid line presents the predation and the dotted line presents the competition.

For the terrestrial ecosystem (Fig. 1a), the interaction network presents the fig-fig wasp mutualism [Compton & Robertson, 1988], which includes four trophic levels: plant (seeds), seed predators (pollinating wasp and non-pollinating wasp), parasitoid wasp and ant. The parasitoid wasp is a specialist predator and only feeds on the pollinating wasp, while the ant is a generalist predator and feeds on all fig wasps. Because the generation span of the fig tree is much longer than both the fig wasps and the ant, we select a new subsystem, which includes pollinating wasp (mutualist), non-pollinating wasp (competitor), parasitoid wasp (specialist predator) and ant (generalist predator), to study the mechanisms for species coexistence in the fig-fig wasp system. For the aquatic ecosystem (Fig. 1b), the complex system diagram shows the plankton community structure [Persson et al., 2001; Benincà et al., 2008], which includes nutrients, producers (edible algae, filamentous algae; bacteria), consumer (heterotrophic nanoflagellates) and zooplankton (daphnia, small grazers). The heterotrophic nanoflagellates species is a specialist predator and only feeds on bacteria, while zooplankton is a generalist predator and feeds on both algae, bacteria and heterotrophic nanoflagellates. In the plankton community, there are two omnivorous links between bacteria and zooplankton. Because nutrients belong to the abiotic factor, which can be considered as an exogenous variable, we also select a new subsystem, which includes bacteria (mutualist), algae (competitor), heterotrophic nanoflagellates (specialist predator) and zooplankton (generalist predator), to study the mechanisms for species coexistence in the plankton community.

In comparing the fig-fig wasp system with the plankton community, we can obtain the similar model framework, which is presented in figure 1c. The new framework includes three trophic levels: generalist predator (*P*_*G*_), specialist predator (*P*_*S*_), mutualist (*N*_1_) and competitor (*N*_2_). *N*_2_ competes with *N*_1_ for resources and *P*_*S*_ only feeds on *N*_1_, while *P*_*G*_ feeds on both *P*_*S*_, *N*_2_ and *N*_1_. This system has three food chains: *N*_1_-*P*_*S*_-*P*_*G*_, *N*_2_-*P*_*G*_ and *N*_1_-*P*_*G*_ (omnivorous chain). The parameters *w*_1_ and *w*_2_ are predation preference coefficients, the solid lines present the predation and the dotted line presents the competition.

### 2.2. The omnivorous food web model

We extend the earlier theoretical work [McCann & Hastings, 1997] to an omnivorous food web model with a competitor:

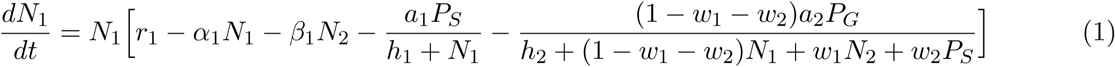

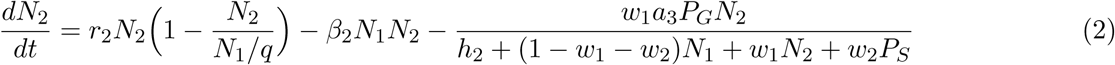

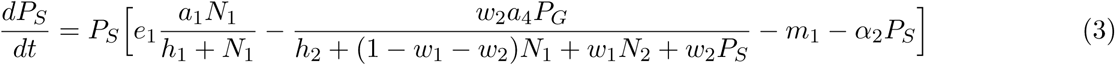

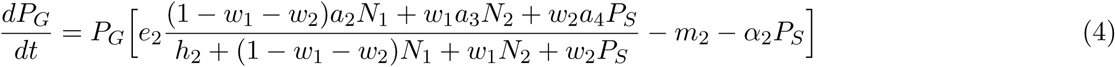

the variable *N*_1_ is the mutualist biomass, *N*_2_ is the competitor biomass, *P*_*S*_ is the biomass of the specialist predator, *P*_*G*_ is the biomass of the generalist predator. The parameters are as follows: *r*_1_ is the growth rate of *N*_1_, *r*_2_ is the growth rate of *N*_2_; *β*_1_ is the interspecific competition coefficient of *N*_2_ to *N*_1_, *β*_2_ is the interspecific competition coefficient of *N*_1_ to *N*_2_; *a*_1_ is the ingestion coefficient of *P*_*S*_ to *N*_1_, *a*_2_ is the ingestion coefficient of *P*_*G*_ to *N*_1_, *a*_3_ is the ingestion coefficient of *P*_*G*_ to *N*_2_, *a*_4_ is the ingestion coefficient of *P*_*G*_ to *P*_*S*_; *h*_1_ and *h*_2_ are the half-saturation densities; *w*_1_ is the predation preference coefficient of *P*_*G*_ to *N*_2_, *w*_2_ is the predation preference coefficient of *P*_*G*_ to *P*_*S*_, 1-*w*_1_-*w*_2_ is the predation preference coefficient of *P*_*G*_ to *N*_1_; *m*_1_ is the mortality of *P*_*S*_, *m*_2_ is the mortality of *P*_*G*_; *e*_1_ is the conversion efficiency of *P*_*S*_, *e*_2_ is the conversion efficiency of *P*_*G*_.

Similar to previous research [Barbier & Loreau, 2019], we induce self-regulation factor into our food webs: *α*_1_ is the density-dependence coefficient of *N*_1_; *α*_2_ is the density-dependence coefficient of *P*_*S*_; *α*_3_ is the density-dependence coefficient of *P*_*G*_. Moreover, we assume the capacity of *N*_2_ is dependent on the development of *N*_1_ and set *N*_1_/*q* acts as the capacity of *N*_2_: when the external resource is rare in the environment, *N*_1_ will produce the resource. For example, the pollinator produces seeds by pollination, and bacteria produces nutrients by decomposition. *q* is a scaling coefficient, when *q* becomes smaller, which means the resource is rich. When *q* becomes larger, this means the resource is rare. The definition of each parameter is presented in Table 1.

**Table 1.** Parameterization of the omnivorous food web model.

| Parameter | Description | Value |
| --- | --- | --- |
| $r_1$ | growth rate of $N_1$ | 0.45 |
| $r_2$ | growth rate of $N_2$ | 0.35 |
| $\alpha_1$ | density-dependence coefficient of $N_1$ | 0.05 |
| $\alpha_2$ | density-dependence coefficient of $P_S$ | 0.02 |
| $\alpha_3$ | density-dependence coefficient of $P_G$ | 0.01 |
| $\beta_1$ | inter-competition coefficient of $N_2$ to $N_1$ | 0.28 |
| $\beta_2$ | inter-competition coefficient of $N_1$ to $N_2$ | 0.1 |
| $a_1$ | ingestion coefficient of $P_S$ to $N_1$ | 0.85 |
| $a_2$ | ingestion coefficient of $P_G$ to $N_1$ | 0.6 |
| $a_3$ | ingestion coefficient of $P_G$ to $N_2$ | 0.3 |
| $a_4$ | ingestion coefficient of $P_G$ to $P_S$ | 0.66 |
| $h_1$ | half-saturation density | 1.4 |
| $h_2$ | half-saturation density | 2.5 |
| $m_1$ | mortality of $P_S$ | 0.3 |
| $m_2$ | mortality of $P_G$ | 0.13 |
| $e_1$ | conversion efficiency of $P_S$ | 1 |
| $e_2$ | conversion efficiency of $P_G$ | 1 |
| $q$ | scaling coefficient | varies in [1.15, 1.6] |
| $w_1$ | predation preference coefficient of $P_G$ to $N_2$ | varies in [0, 1] |
| $w_2$ | predation preference coefficient of $P_G$ to $P_S$ | varies in [0, 1] |
| $1 - w_1 - w_2$ | predation preference coefficient of $P_G$ to $N_1$ | varies in [0, 1] |

Then, similar to earlier work on trophic interactions [McCann et al., 1998], we define interaction strength of predation (*I*) and relative interaction strength (*RIS*):

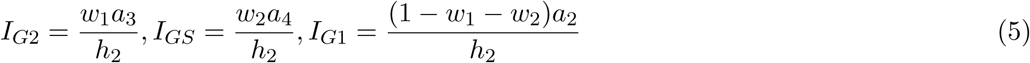

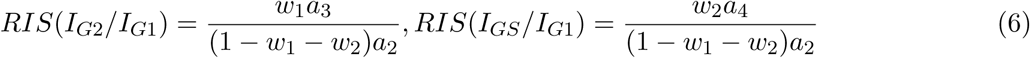

### 2.3. Parameterization and model simulation

Our omnivorous food web belongs to a general model framework, all model parameters (Table 1) can be chosen based on the stability criteria [Temam, 2012] and bifurcation method [Holmes & Guckenheimer, 2002] under the nonnegative equilibrium. First, we calculate a positive equilibrium of species coexistence and analyze the maximum real part of all eigenvalues (Re(*λ*_*max*_)) under the equilibrium, and obtain a stable state when Re(*λ*_*max*_) is negative. Second, based on the bifurcation method (Hopf bifurcation or transcritical bifurcation), we select both stability and species number as assessment indicators: Hopf bifurcation happens when the species number is unchanged and stability changes between stable and unstable states, while transcritical bifurcation emerges when the stability is unchanged and species number changes between increasing and decreasing states.

We regulate the preference coefficients (*w*_1_ and *w*_2_) to look for the parameter space of species coexistence and analyze how food web structure changes with the increase of the relative interaction strength (*RIS*). Moreover, we use a two-parameter bifurcation method to show how both species diversity and system stability change with the interaction strength under the different resource conditions, and study the robust of our model results by using the Holling I function response in Appendix A. Finally, we combine theoretical predictions with empirical data analyses. All simulations are carried out using the Matlab 7.0 [MathWorks, 2005] implementation.

## 3. Results

### 3.1. Population dynamic simulation

Model simulations show that both predation preference coefficients (*w*_1_ and *w*_2_) and resource scaling coefficient (*q*) will change food web structure and system stability (Fig. 2). First, in a low resource level (*q*=1.45), we show that food web complexity changes with both *w*_1_ and *w*_2_: stable coexistence of species (*w*_1_=0.1, *w*_2_=0.2; Fig. 2a) and oscillatory coexistence of species (*w*_1_=0.3, *w*_2_=0.25; Fig. 2b) in our food web model; stable coexistence (*w*_1_=0.34, *w*_2_=0.6; Fig. 2e) and oscillatory coexistence (*w*_1_=0.11, *w*_2_=0.86; Fig. 2f) of both *N*_1_, *P*_*S*_ and *P*_*G*_; stable coexistence of both *N*_1_, *N*_2_ and *P*_*G*_ (*w*_1_=0.1, *w*_2_=0.4; Fig. 2c); stable coexistence of both *N*_1_ and *P*_*G*_ (*w*_1_=0.15, *w*_2_=0.6; Fig. 2g); oscillatory coexistence of both *N*_1_, *P*_*S*_ and *N*_2_ (*w*_1_=0.8, *w*_2_=0.1; Fig. 2i). Second, in an intermediate resource level (*q*=1.28), we found that stable coexistence of both *N*_1_, *P*_*S*_ and *N*_2_ can be obtained (*w*_1_=0.3, *w*_2_=0.43; Fig. 2h). Third, oscillatory coexistence of both *N*_1_, *N*_2_ and *P*_*G*_ can be maintained (*w*_1_=0, *w*_2_=0.66; Fig. 2d) under a high resource level (*q*=1.15). Moreover, we found that simulating the chain ‘*N*_1_-*P*_*G*_’ cannot result in an oscillatory dynamics, instead of a new stable structure (such as structure in figure 2c or figure 2e).

**Fig. 2.**
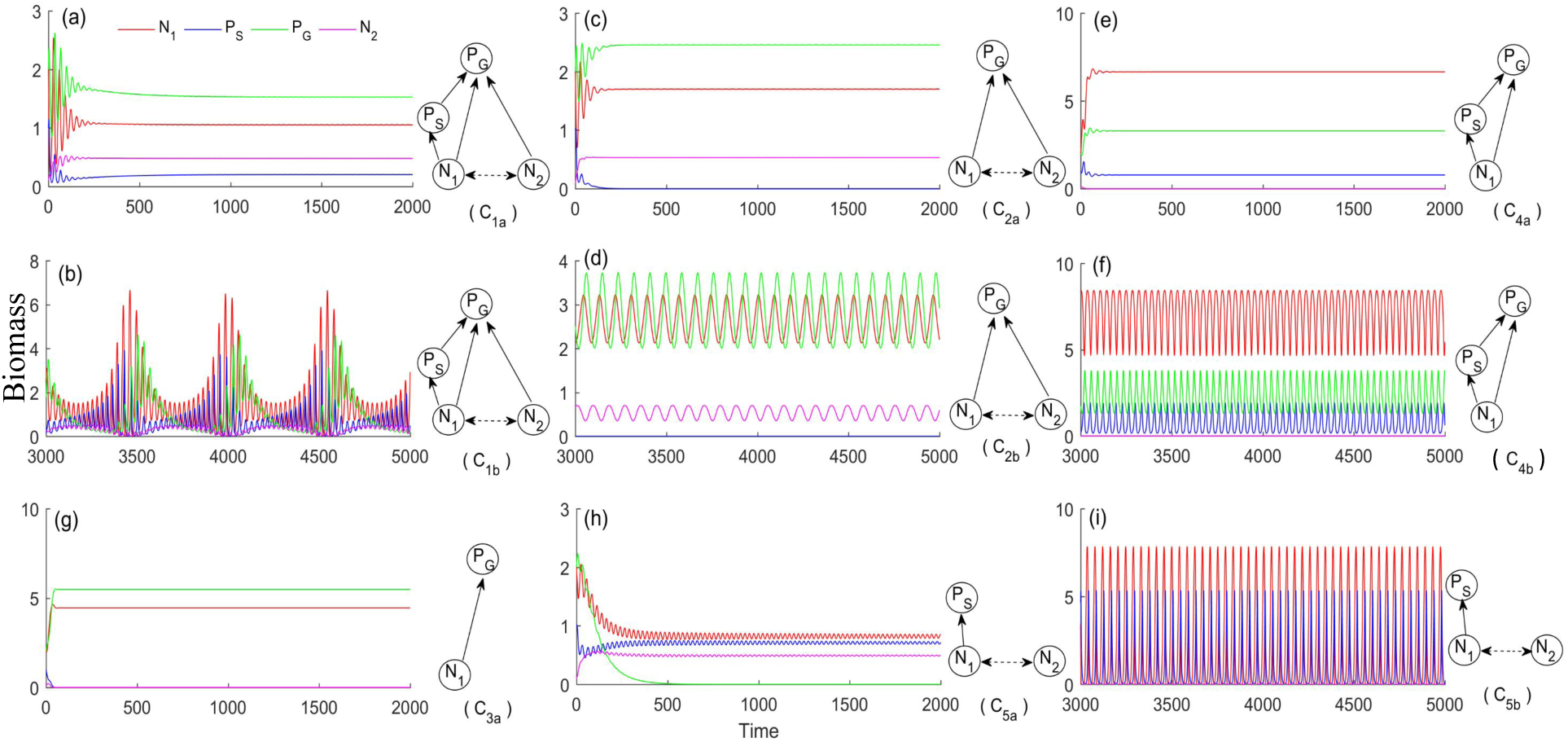
Population biomass changes with predation preference coefficients and resource scaling coefficient. (a) *q*=1.45, *w*_1_=0.1, *w*_2_=0.2; (b) *q*=1.45, *w*_1_=0.3, *w*_2_=0.25; (c) *q*=1.45, *w*_1_=0.1, *w*_2_=0.4; (d) *q*=1.15, *w*_1_=0, *w*_2_=0.66; (e) *q*=1.45, *w*_1_=0.34, *w*_2_=0.6; (f) *q*=1.45, *w*_1_=0.11, *w*_2_=0.86; (g) *q*=1.45, *w*_1_=0.15, *w*_2_=0.6; (h) *q*=1.28, *w*_1_=0.3, *w*_2_=0.43; (i) *q*=1.45, *w*_1_=0.8, *w*_2_=0.1. *C*_1*a*_means structure *C*_1_ has a stable dynamics, while *C*_1*b*_ means structure *C*_1_ produces an oscillatory dynamics. All parameters are presented in Table 1.

### 3.2. Relative interaction strength changes species coexistence

Next, in the fixed resource level (*q*=1.45; Fig. 2), we find that the population dynamics and food web structure change with the increase of the relative interaction strength *RIS*(*I*_*G*2_/*I*_*G*1_) (Fig. 3) and *RIS*(*I*_*GS*_/*I*_*G*1_) (Fig. 4).

**Fig. 3.**
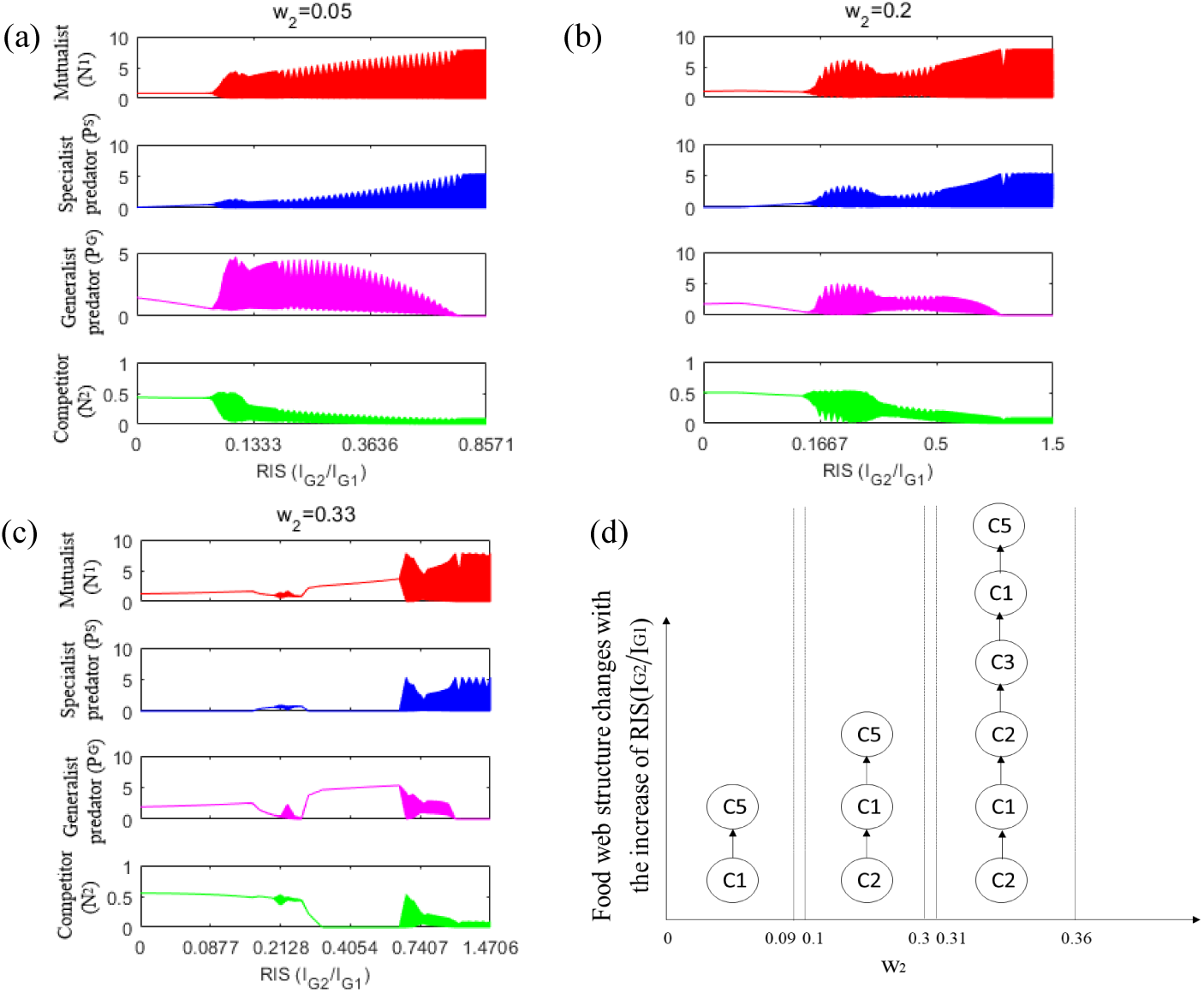
Population biomass and food web structure change with *RIS*(*I*_*G*2_/*I*_*G*1_) under the different predation preference *w*_2_. (a-c) bifurcation diagrams; (d) evolutionary trend of food web structure. All parameters are presented in Table 1 except for *q*=1.45.

**Fig. 4.**
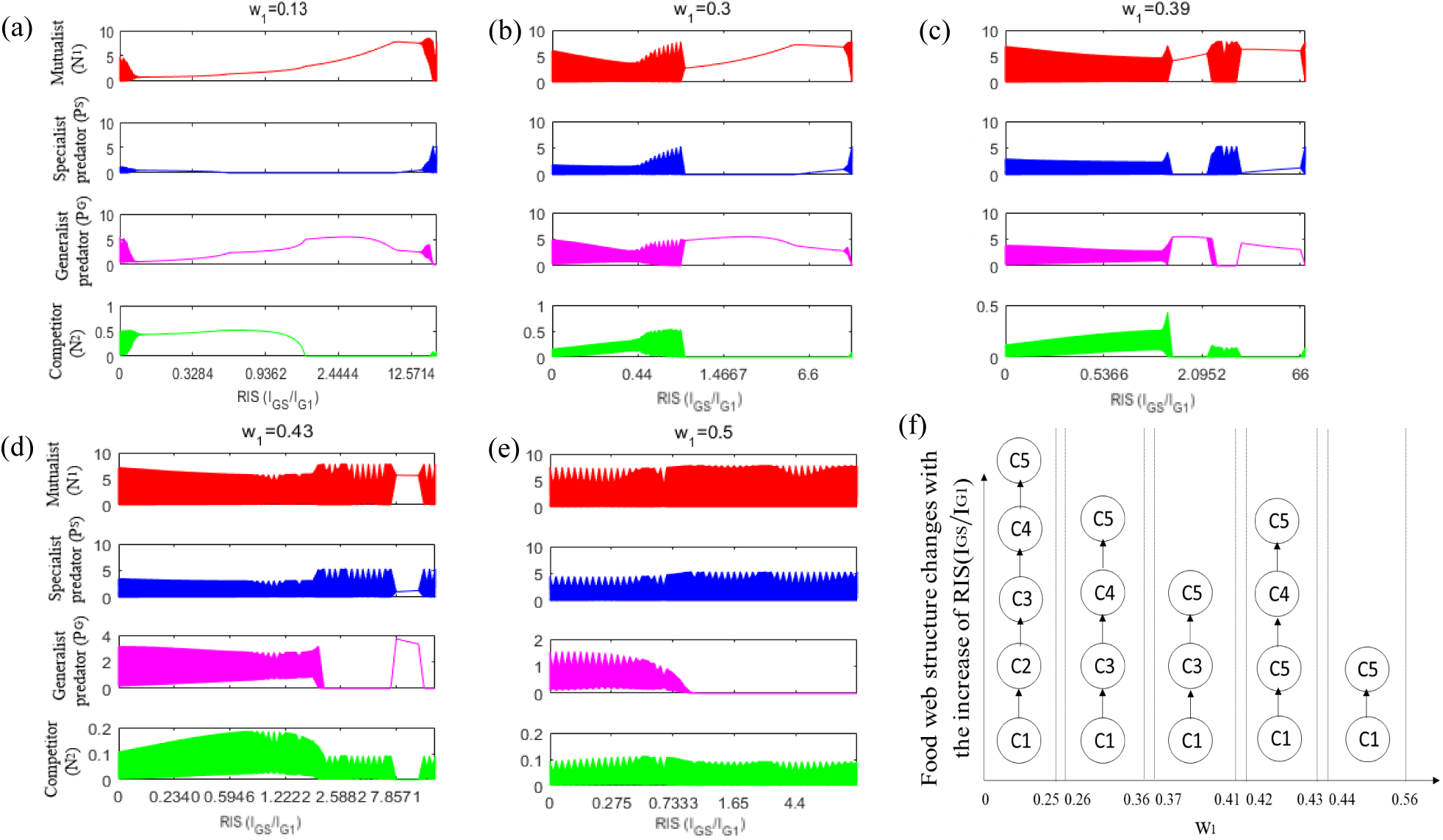
Population biomass and food web structure change with *RIS*(*I*_*GS*_ /*I*_*G*1_) under the different predation preference *w*_1_. (a-e) bifurcation diagrams; (f) evolutionary trend of food web structure. All parameters are presented in Table 1 except for *q*=1.45.

On one hand, we find there is no coexistence of species with changing *w*_1_ when *w*_2_ ∈ [0.37, 1]. The coexistence of species can be obtained when *w*_2_ ∈ [0, 0.36] (Fig. 3). When the predation preference of *P*_*G*_ to *P*_*S*_ is at a low strength (*w*_2_ ∈ [0, 0.09]), let *w*_2_=0.05, and with increasing *RIS*(*I*_*G*2_/*I*_*G*1_), first, species coexistence, then *P*_*G*_ goes extinct (Fig. 3a). Simulating the model with any *w*_2_ (*w*_2_ ∈ [0, 0.09]), we can obtain the same process of ‘species coexistence–*P*_*G*_ extinction’. When *P*_*G*_ has an intermediate predation preference for *P*_*S*_ (*w*_2_ ∈ [0.1, 0.3]), let *w*_2_=0.2, and with increasing *RIS*(*I*_*G*2_/*I*_*G*1_), first, *P*_*S*_ goes extinct, then species coexistence, and finally, *P*_*G*_ goes extinct (Fig. 3b). Simulating the model with any *w*_2_ (*w*_2_∈ [0.1, 0.3]), we can obtain the same process of ‘*P*_*S*_ extinction–species coexistence–*P*_*G*_ extinction’. When *P*_*G*_ has a high predation preference (*w*_2_ ∈ [0.31, 0.36]), let *w*_2_=0.33, and with increasing *RIS*(*I*_*G*2_/*I*_*G*1_), first, *P*_*S*_ goes extinct, then species coexistence, *P*_*S*_ goes extinct again, both *P*_*S*_ and *N*_2_ go extinct, species coexistence, and finally, *P*_*G*_ goes extinct (Fig. 3c). Simulating the model with any *w*_2_ (*w*_2_ ∈ [0.31, 0.36]), we can obtain the same process of ‘*P*_*S*_ extinction–species coexistence–*P*_*S*_ extinction–*P*_*S*_, *N*_2_ extinction–species coexistence–*P*_*G*_ extinction’. Finally, we can obtain the evolutionary trend of food web structure changes with the increase of *RIS*(*I*_*G*2_/*I*_*G*1_) under the different predation preference *w*_2_ (Fig. 3d).

On the other hand, we also find there is no coexistence of species with changing *w*_2_ when *w*_1_ ∈ [0.57, 1]. The coexistence of species can be obtained when *w*_1_ ∈ [0, 0.56] (Fig. 4). When the predation preference of *P*_*G*_ to *N*_2_ is at a lower strength (*w*_1_ ∈ [0, 0.25]), let *w*_1_=0.13, and with increasing *RIS*(*I*_*GS*_/*I*_*G*1_), first, species coexistence, then *P*_*S*_ goes extinct, both *P*_*S*_ and *N*_2_ go extinct, *N*_2_ goes extinct, and finally, *P*_*G*_ goes extinct (Fig. 4a). Simulating the model with any *w*_1_ (*w*_1_ ∈ [0, 0.25]), we can obtain the same process of ‘species coexistence–*P*_*S*_ extinction–*P*_*S*_, *N*_2_ extinction–*N*_2_ extinction–*P*_*G*_ extinction’. When *P*_*G*_ has a low predation preference for *N*_2_ (*w*_1_ ∈ [0.26, 0.36]), let *w*_1_=0.3, and with increasing *RIS*(*I*_*GS*_/*I*_*G*1_), first, species coexistence, then both *P*_*S*_ and *N*_2_ go extinct, *N*_2_ goes extinct, and finally, *P*_*G*_ goes extinct (Fig. 4b). Simulating the model with any *w*_1_ (*w*_1_ ∈ [0.26, 0.36]), we can obtain the same process of ‘species coexistence–*P*_*S*_, *N*_2_ extinction–*N*_2_ extinction–*P*_*G*_ extinction’. When *P*_*G*_ has an intermediate predation preference (*w*_1_ ∈ [0.37, 0.41]), let *w*_1_=0.39, and with increasing *RIS*(*I*_*GS*_/*I*_*G*1_), first, species coexistence, then both *P*_*S*_ and *N*_2_ go extinct, *P*_*G*_ goes extinct, *N*_2_ goes extinct, and finally, *P*_*G*_ goes extinct again (Fig. 4c). Simulating the model with any *w*_1_ (*w*_1_ ∈ [0.37, 0.41]), we can obtain the same process of ‘species coexistence–*P*_*S*_, *N*_2_ extinction–*P*_*G*_ extinction–*N*_2_ extinction–*P*_*G*_ extinction’. When *P*_*G*_ has a high predation preference (*w*_1_ ∈ [0.42, 0.43]), let *w*_1_=0.43, and with increasing *RIS*(*I*_*GS*_/*I*_*G*1_), first, species coexistence, then *P*_*G*_ goes extinct, *N*_2_ goes extinct, and finally, *P*_*G*_ goes extinct again (Fig. 4d). Simulating the model with any *w*_1_ (*w*_1_ ∈ [0.42, 0.43]), we can obtain the same process of ‘species coexistence–*P*_*G*_ extinction–*N*_2_ extinction–*P*_*G*_ extinction’. When *P*_*G*_ has a higher predation preference (*w*_1_ ∈ [0.44, 0.56]), let *w*_1_=0.5, and with increasing *RIS*(*I*_*GS*_/*I*_*G*1_), first, species coexistence, then *P*_*G*_ goes extinct (Fig. 4e). Simulating the model with any *w*_1_ (*w*_1_ ∈ [0.44, 0.56]), we can obtain the same process of ‘species coexistence–*P*_*G*_ extinction’. Finally, we can also obtain the evolutionary trend of food web structure changes with the increase of *RIS*(*I*_*GS*_/*I*_*G*1_) under the different predation preference *w*_1_ (Fig. 4f). By comparing these results (Figs. 3, 4), we find that evolutionary trend of food web structure changes with the relative predation strength (Fig. 4) is more diversity than the relative competition strength (Fig. 3).

### 3.3. Interaction strength regulates food web structure and system stability

Different with the single-parameter bifurcation analyses (Figs. 3, 4), a two-parameter bifurcation approach shows that both food web structure and system stability can change with the interaction strength (*I*_*G*2_ and *I*_*GS*_) under the different resource conditions (scaling coefficient *q*) (Fig. 5).

**Fig. 5.**
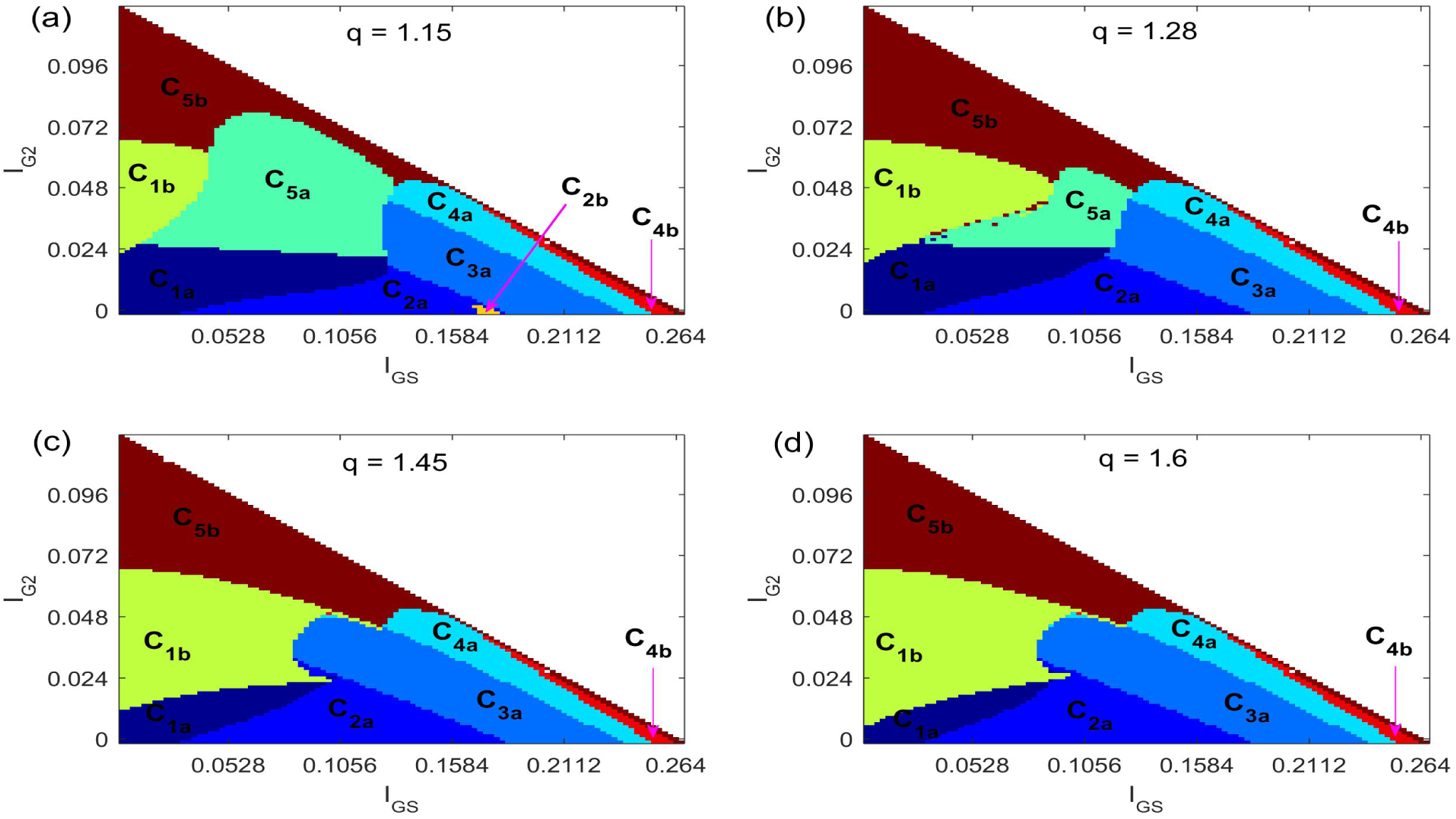
Interaction strength regulates food web structure and system stability under the different resource levels. All parameters are presented in Table 1.

For one thing, based on a crosswise contrast approach of the fixed resource level (*q*), we can find three meaningful results. First, weak interaction strength (*I*_*G*2_ < 0.024, *I*_*GS*_ < 0.1056) promotes a stable coexistence of species (region *C*_1*a*_) in our omnivorous food web model. Second, weak to intermediate interaction strength (*I*_*G*2_ < 0.072, *I*_*GS*_ < 0.1056) promotes an oscillatory coexistence of species (region *C*_1*b*_). Third, increasing the interaction strength (*I*_*G*2_) between *P*_*G*_ and *N*_2_ will result in Hopf bifurcation, that is, system stability changes and food web structure is unchanged (regions *C*_1*a*_ and *C*_1*b*_; regions *C*_5*a*_ and *C*_5*b*_); while increasing the interaction strength (*I*_*GS*_) between *P*_*G*_ and *P*_*S*_ will cause transcritical bifurcation, that is, system stability is unchanged and food web structure changes between increasing and decreasing states (regions *C*_1*a*_ and *C*_2*a*_; regions *C*_2*a*_ and *C*_3*a*_; regions *C*_3*a*_ and *C*_4*a*_).

For another, by adopting a longitudinal contrast approach of the different resource levels (*q*), we obtain some important results. First, increasing the scaling coefficient *q* (less resource in food webs) will shrink the region of stable coexistence of species (*C*_1*a*_) and expand the region of oscillatory coexistence of species (*C*_1*b*_). For example, we can get the largest region of the stable coexistence of species (*C*_1*a*_) under the highest resource level (*q*=1.15; Fig. 5a) than in other three cases (Fig. 5b-d), and obtain the largest region of the oscillatory coexistence of species (*C*_1*b*_) under the lowest resource level (*q*=1.6; Fig. 5d) than in other three cases (Fig. 5a-c). Second, decreasing the scaling coefficient *q* (more resource in food webs) will increase the stability types of food web structure. For example, when considering a lower resource level (Fig. 5c-d), we find that only seven regions emerge (*C*_1*a*_, *C*_1*b*_, *C*_2*a*_, *C*_3*a*_, *C*_4*a*_, *C*_4*b*_ and *C*_5*b*_); under a higher resource level (Fig. 5a-b), some new regions emerge in figure 5a (*q*=1.15; *C*_2*b*_ and *C*_5*a*_) and figure 5b (*q*=1.28; *C*_5*a*_). Third, comparing with the different resource levels (Fig. 5a-d), we find the regions (*C*_5*a*_ and *C*_5*b*_) of *P*_*G*_ extinction will shrink with the decrease of resource (the increased scaling coefficient *q*) in food webs; moreover, each diagonal line falls into region *C*_5*b*_, which means *P*_*G*_ will always become extinct if *P*_*G*_ does not prey on *N*_1_ (*w*_1_+*w*_2_ = 1; Fig. 1c). Finally, we obtain the similar results by selecting the Holling I function response in the omnivorous food web model (Appendix A).

### 3.4. Comparison between theoretical predictions and empirical data

In this section, we compare model simulation with empirical data. First, we set *w*_1_=0.33 (*I*_*G*2_=0.0396), *w*_2_=0.25 (*I*_*GS*_=0.066), only vary *q* (*q* ∈ [1.28, 1.6]; Fig. 5b-d), model simulations show that an oscillatory coexistence of species will emerge (*C*_1*b*_ in figure 5b-d). Then, we can obtain the bifurcation diagrams of four species biomasses (Fig. 6a-d).

**Fig. 6.**
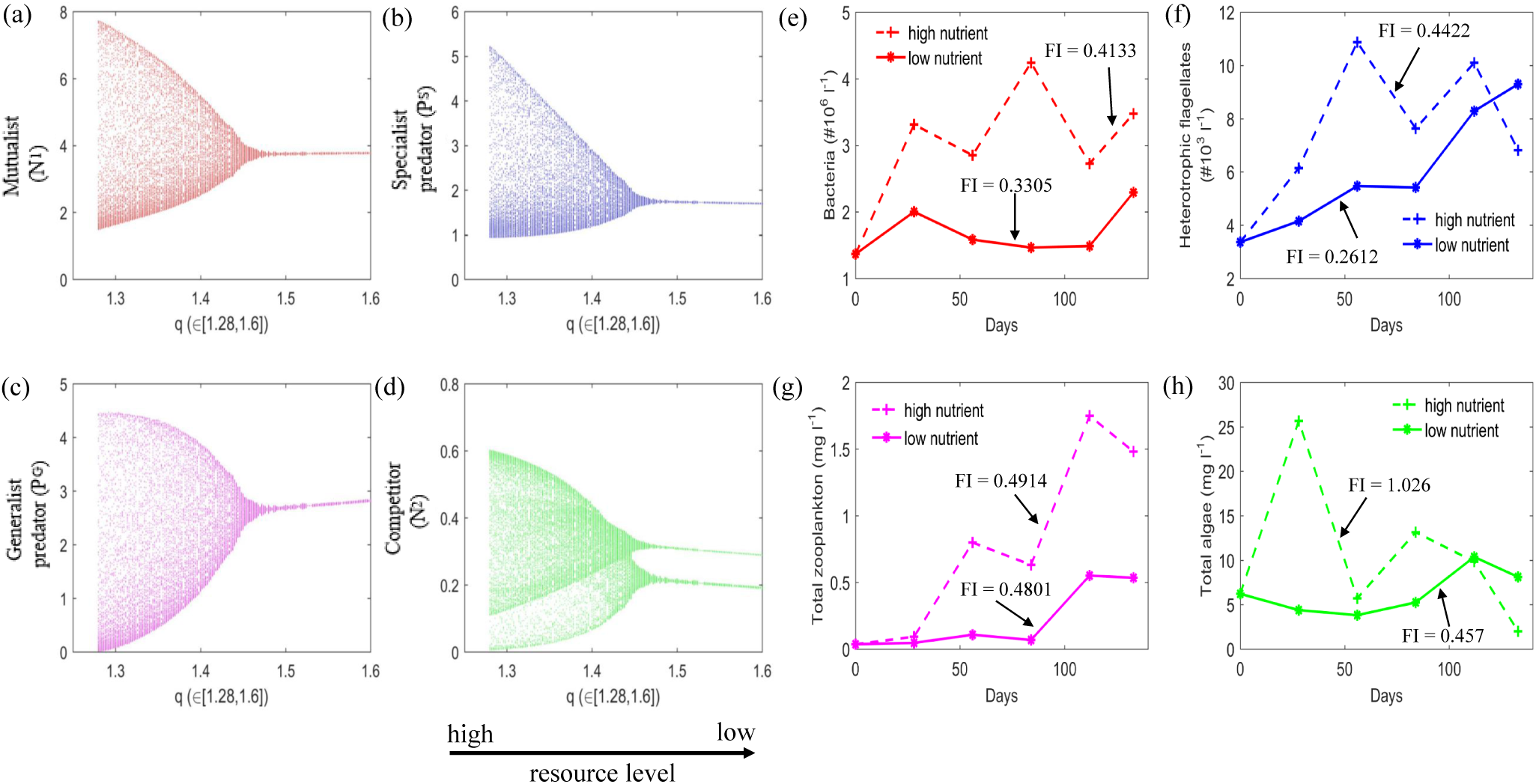
Comparison between theoretical prediction and empirical data. (a-d) Population biomass changes with the scaling coefficient *q*. (e-h) Population abundance/biomass over time in experimental tanks received either high (dotted line marked in ‘+’) or low (solid line marked in ‘*’) nutrient additions (Persson et al., 2001). All parameters are presented in Table 1 except for *w*_1_=0.33, *w*_2_=0.25 and *q* ∈ [1.28, 1.6].

The model simulations show that magnitude of each biomass will continuously decrease with the increase of the scaling coefficient *q*. When *q* becomes larger, this means the resource level is at a lower condition. So the magnitude under a rich resource (*q*=1.28, irregular/chaotic oscillation) is higher than the poor (*q*=1.6, regular/periodic oscillation). This result is consistent with the previous empirical data (Fig. 6e-h), which is presented in empirical results of the food web II [Persson et al., 2001]. By comparing high (dotted line marked in ‘+’) nutrient level with low (solid line marked in ‘*’) nutrient level, we calculate fluctuation index (*FI*) of a series [Dey & Joshi, 2006] and find that the magnitude of both bacteria (*N*_1_; Fig. 6e), heterotrophic nanoflagellates (*P*_*S*_; Fig. 6f), total zooplankton (*P*_*G*_; Fig. 6g) and total algae (*N*_2_; Fig. 6h) under the high nutrient state is larger than the magnitude under the low nutrient state: in figure 6e, *FI*_*high*_ = 0.4133, *FI*_*low*_ = 0.3305; in figure 6f, *FI*_*high*_ = 0.4422, *FI*_*low*_ = 0.2612; in figure 6g, *FI*_*high*_ = 0.4914, *FI*_*low*_ = 0.4801; in figure 6h, *FI*_*high*_ = 1.026, *FI*_*low*_ = 0.457. So, our theoretical results can predict the population oscillation dynamics of empirical data.

## 4. Discussion

Species coexistence makes ecologists fascinated [Gravel et al., 2011; Levine et al., 2017], and the mechanisms of species coexistence have been extensively discussed in a variety of ecosystems, such as higher-order interaction [Bairey et al., 2016; Grilli et al., 2017], competition interactions [Foster & Bell, 2012; Vandermeer et al., 2002], predation interactions [Abrams, 1999; Ishii & Shimada, 2012] and chaotic oscillation [Allen et al., 1993; Károlyi et al., 2000]. However, there is still a lack of the general mechanisms of species coexistence in the real ecosystems. Here we establish a general omnivorous food web model with a competitor. We analyze species diversity and food web stability by regulating the interaction strength of the generalist predator (top predator) to both mutualist, competitor and specialist predator, and find the generalist predator can regulate species coexistence and system stability under the different resource/nutrition levels. Moreover, the model results can qualitatively predict the oscillation of empirical data, that is, a decrease in resource/nutrient supply may result in an increase of stability (such as transition from regular periodic oscillation to irregular/chaotic oscillation), which is consistent with the result of ‘paradox of enrichment’ [Rosenzweig, 1971]. Our theoretical predictions show that weak predation strength can promote stable coexistence of predators and prey, which is consistent with the newest empirical research: weak interactions are highly correlated with a higher biodiversity in a natural fish community [Ushio et al., 2018].

### 4.1. Omnivory facilitates population dynamics of top predator

In our model simulation, when omnivorous link does not exist in the food web model, that is, *w*_1_+*w*_2_ = 1 (the diagonal of region *C*_5*b*_), the top predator (generalist predator *P*_*G*_) will always become extinct. To slightly increase the omnivory means to increase the long connection in our system, which makes food web structure more stable (*C*_5*a*_, *C*_4*a*_ and *C*_3*a*_). This result is consistent with earlier theoretical work [Fussmann & Heber, 2002]: increasing the number of omnivorous links will reduce the probability of chaos in food web models, which means the system will become more stable. Moreover, our omnivorous food web model is derived from the previous model [McCann & Hastings, 1997], which showed that the addition of omnivory locally stabilizes the food web in a very complete way. Although the parameter sets of the two models are significantly different, their conclusions are similar, that is, omnivory can promote food web stability.

### 4.2. Increased competition promotes species coexistence

We add a competitor into the general food web structure [McCann & Hastings, 1997], and find that the new food web model has a rich dynamical behaviour, such as the interaction between competition and species coexistence. Our theoretical analyses show that the increased competition can promote species coexistence in the omnivorous food webs. In our system, the interaction strength *I*_*G*2_ indicates the increased competition of *N*_2_ to *N*_1_: the competition is a higher level when the predation pressure of *P*_*G*_ to *N*_2_ (*I*_*G*2_) has a lower value. For one thing, a strong competition promotes stable coexistence of species (*C*_1*a*_). For another, an intermediate to strong competition facilitates an oscillatory coexistence of species (*C*_1*b*_). Our results are consistent with earlier theoretical study [Vandermeer et al., 2002], which shows that the increased competition promotes species coexistence in a theoretical framework. Moreover. our results also can support earlier empirical work [Foster & Bell, 2012], which shows that the competition dominates interactions among culturable microbial species.

### 4.3. Top-down control changes food web structure and system stability

In the omnivorous food web, the top-down effects are represented by the interaction strength (*I*_*G*2_ and *I*_*GS*_). Our theoretical predictions show that the weak top-down control (lower values of *I*_*G*2_ and *I*_*GS*_) can promote stable coexistence of species (*C*_1*a*_), while the weak to intermediate top-down control (moderate values of *I*_*G*2_ and *I*_*GS*_) can promote an oscillatory coexistence of species (*C*_1*b*_). However, a strong top-down control (higher values of *I*_*G*2_ and *I*_*GS*_) will change food web structure and stability (*C*_3*a*_, *C*_4*a*_, *C*_4*b*_ and *C*_5*b*_). Our results are consistent with earlier work [McCann et al., 1998], which maintains that weak to intermediate strength links are important in promoting food web persistence and stability. Our model results also support both theoretical [Abrams, 1999] and empirical work [Snyder & Ives, 2003; Ishii & Shimada, 2012], which show that the predator can mediate coexistence of species. Moreover, the newest experimental work showed that top-down control of top predator will change niche structure and species coexistence [Pringle et al., 2019]. Our theoretical predictions are also consistent with the newest experimental study on the fig-fig wasp mutualism [Wang et al., 2019], which showed that loss of top-down control (parasitoidism on the pollinators) resulted in a change in the balance of reciprocal benefits, thereby changing food web complexity. In our work, changing predation preference strength will affect the outcomes of top-down control in the food webs, thereby regulate food web structure and system stability.

### 4.4. Bottom-up control promotes species diversity and food web stability

In our model, we introduce a key parameter *q*, which can indicate the resource/nutrition level: a higher scaling coefficient *q* indicates a lower resource/nutrition condition. Our theoretical results find two important results: first, a lower resource/nutrition level (a higher value of *q*) results in a larger region of species coexistence (both *C*_1*a*_ and *C*_1*b*_); second, as the decrease of resource/nutrition level, the magnitude of population oscillation will shrink, such as oscillation magnitude decreases from irregular/chaotic oscillation to regular/periodic oscillation. These results show that enrichment can lead to a change in dynamic states, which is the bases of bottom-up control in the community [Hairston et al., 1960]. In the fig-fig wasp system, the host tree will reduce the nutrition resources to the fig with less pollen or without pollen, and this will result in a lower rate of reproductive success for all the wasps in the fig [Wang et al., 2010]. Our model predictions are consistent with the previous work [Moore & de Ruiter, 2012; Rosenzweig, 1971], which show that resource limitations have a stabilizing effect on systems. In our study, bottom-up control under a low resource condition can promote system stability and facilitate species coexistence (oscillatory dynamics).

### 4.5. Conclusions

In this work, we add a competitor of the mutualist in an omnivorous food web to re-evaluate the general mechanisms for species coexistence, the results show that weak predation strength can promote stable coexistence of predators and prey. Moreover, evolutionary trend of food web structure changes with the relative predation strength is more diversity than the relative competition strength, and an integration of both omnivory, increased competition, top-down control and bottom-up control can promote species coexistence. Our theoretical work could enrich the general omnivorous theory on species coexistence and system stability in the real food webs, and may help understand the interactions among omnivory, food web stability and species diversity.

## Acknowledgments

This work was supported by funds from the Exchange Funding of Northwestern Polytechnical University to L.W.. We thank Gabriel Gellner and Jennifer Meunier for helpful comments on the manuscript. R.W.W. conceived the study and all authors participated in study design. L.W. carried out analyses and wrote the first draft of the paper. All authors contributed substantially to revisions.

## Appendix A

We use the Holling I function response in the omnivorous food web model with a competitor:

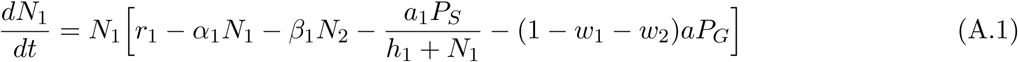

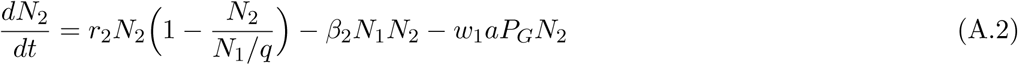

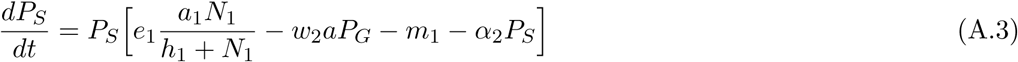

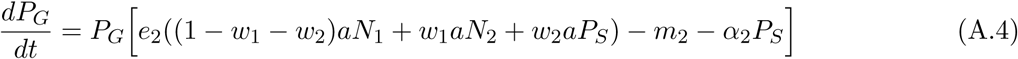

the new parameter *a* is the mean attack rate of *P*_*G*_ to its prey. Then, we use the two-parameter bifurcation method to show how both species diversity and system stability change with the interaction strength under the different resource conditions. Comparing with the case of Holling II (Fig. 5), we obtain the similar results by selecting the Holling I function response in the omnivorous food web model (Fig. A.1).

**Fig. A.1.**
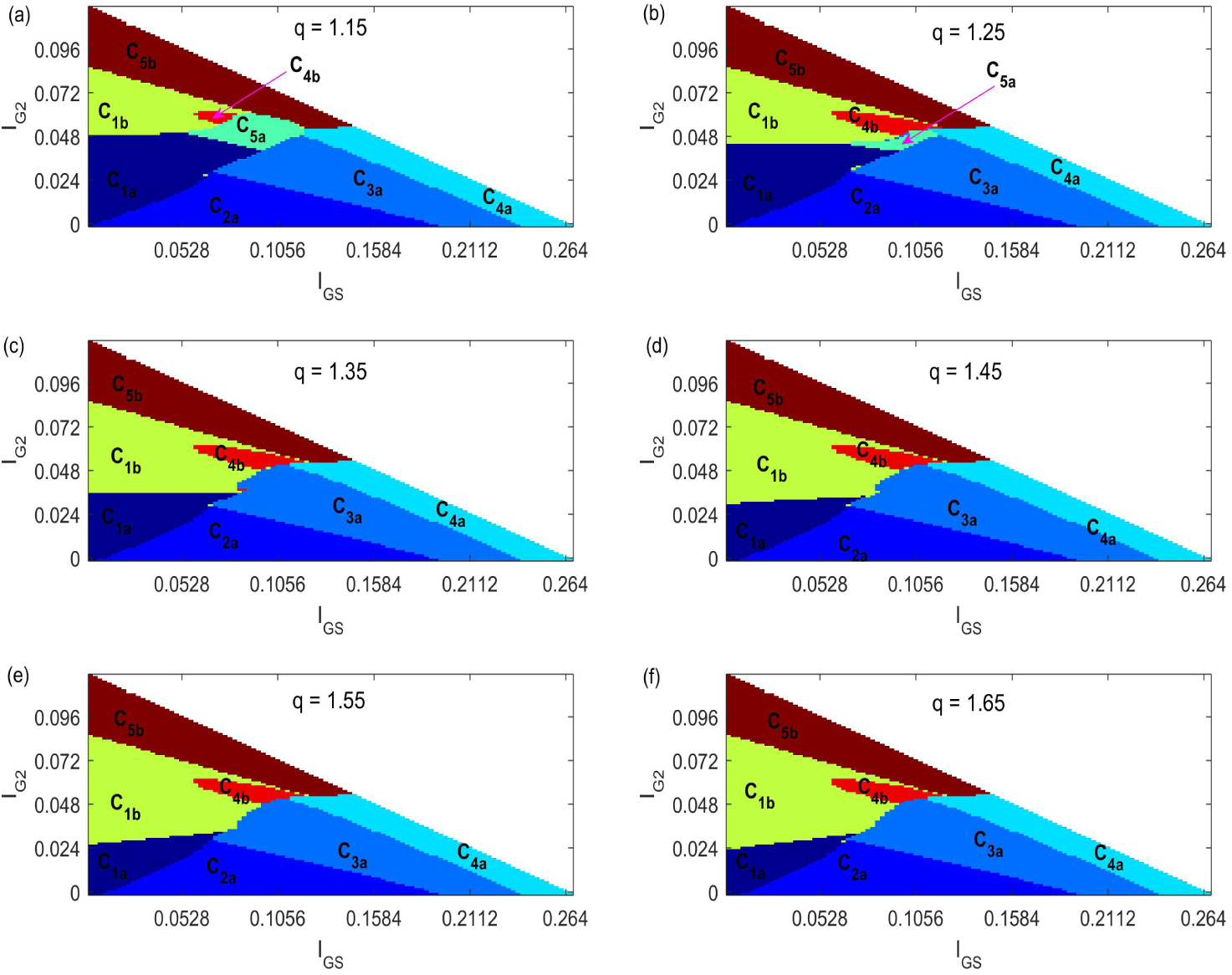
Interaction strength regulates food web structure and system stability under the different resource levels. All parameters are presented in Table 1 except for *a*=0.19.

Based on a crosswise contrast approach of the fixed resource level (*q*), we can find some similar results. First, weak interaction strength (lower *I*_*GS*_ and *I*_*G*2_) promotes a stable coexistence of species (region *C*_1*a*_). Second, weak to intermediate interaction strength (moderate *I*_*G*2_ and *I*_*GS*_) promotes an oscillatory coexistence of species (region *C*_1*b*_). Third, increasing the interaction strength (*I*_*G*2_) between *P*_*G*_ and *N*_2_ mainly result in Hopf bifurcation, that is, system stability changes and food web structure is unchanged (regions *C*_1*a*_ and *C*_1*b*_ in figure A.1a-f; regions *C*_5*a*_ and *C*_5*b*_ in figure A.1a); while increasing the interaction strength (*I*_*GS*_) between *P*_*G*_ and *P*_*S*_ mainly result in transcritical bifurcation, that is, system stability is unchanged and food web structure changes between increasing and decreasing states (regions *C*_1*a*_ and *C*_2*a*_; regions *C*_2*a*_ and *C*_3*a*_; regions *C*_3*a*_ and *C*_4*a*_).

Then, by adopting a longitudinal contrast approach of the different resource levels (*q*), we obtain some important results. For one thing, increasing the scaling coefficient *q* (less resource in food webs) will shrink the region of stable coexistence of species (region *C*_1*a*_) and expand the region of oscillatory coexistence of species (region *C*_1*b*_). For example, we can get the largest region of the stable coexistence of species (region *C*_1*a*_) under the highest resource level (*q*=1.15; Fig. A.1a) than other five cases (Fig. A.1b-f), and the largest region of the oscillatory coexistence of species (region *C*_1*b*_) under the lowest resource level (*q*=1.65; Fig. A.1f) than other five cases (Fig. A.1a-e). For another, decreasing the scaling coefficient *q* (more resource in food webs) will increase the stability types of food web structure. For example, when considering a lower resource level (Fig. A.1c-f), we find that only seven regions emerge (regions *C*_1*a*_, *C*_1*b*_, *C*_2*a*_, *C*_3*a*_, *C*_4*a*_, *C*_4*b*_ and *C*_5*b*_); under a higher resource level (Fig. A.1a-b), a new region (region *C*_5*a*_) emerges. Finally, comparing with the different resource levels, we find the regions (regions *C*_5*a*_ and *C*_5*b*_) of *P*_*G*_ extinction will shrink with the decrease of resource (the increased scaling coefficient *q*) in food webs.

